# Programmed DNA elimination drives rapid genomic innovation in two thirds of all bird species

**DOI:** 10.1101/2025.07.16.664580

**Authors:** Francisco J. Ruiz-Ruano, Stephen A. Schlebusch, Niki Vontzou, Hugo Moreno, Matthew T. Biegler, Verena E. Kutschera, Diana Ekman, Inês Borges, Yifan Pei, Roberto Rossini, Tomas Albrecht, Jesper Boman, Pavel Borodin, Reto Burri, Kristal E. Cain, Wolfgang Forstmeier, Carolina Frankl-Vilches, Manfred Gahr, Simon C. Griffith, Amy M. Hill, Martin Irestedt, Leo Joseph, Knud A. Jønsson, Takeshi Kawakami, Bart Kempenaers, Lyubov Malinovskaya, Jakob C. Mueller, Edivaldo H. C. de Oliveira, Octavio M. Palacios-Gimenez, Vaidas Palinauskas, Anna Qvarnström, Radka Reifova, Jakub Ridl, J. Carolina Segami, David J. X. Tan, Anna Torgasheva, Annabel Whibley, Alexander Suh

## Abstract

Bird genomes are among the most stable in terms of synteny and gene content across vertebrates. However, germline-restricted chromosomes (GRCs) represent a striking exception where programmed DNA elimination confines large-scale genomic changes to the germline. GRCs are known to occur in songbirds (oscines), but have been studied only in a few species of Passerides such as the zebra finch, the key model for passerine genomics. Their presence and evolutionary dynamics in most major passerine lineages remain largely unexplored, with suboscines entirely unexamined by cytogenetic or genomic methods. Here, we present the most comprehensive comparative analysis of GRCs to date, spanning 44 million years of passerine evolution. By generating the first germline reference genomes of an oscine and a suboscine, 22 novel germline draft genomes spanning nearly all major passerine lineages and a germline draft genome of a parrot outgroup, we show that the GRC is likely present in 6,700 passerine species. Our results reveal that the GRC evolves rapidly and distinctly from the standard A chromosomes (autosomes and sex chromosomes), yet retains functionally important, selectively maintained genes. We observed gene and repeat turnover occuring orders of magnitude faster than on the A chromosomes. Some GRC genes, such as *cpeb1* and *pim1*, are widespread from an ancient duplication. In contrast, other GRC genes, like *mfsd2b* and *bmp15*, have been independently duplicated onto the GRC multiple times, suggesting adaptive constraints. The discovery of *zglp1* on the zebra finch GRC, initially copied from chromosome 30 and subsequently lost from it, indicates functional replacement, where the GRC permits gene loss from the standard genome. As the GRC harbors the only *zglp1* copy in most of the ∼4000 Passerides species, GRC loss would compromise essential germline functions. Our findings establish the GRC as a genomic innovator driving rapid germline evolution. This fact highlights its evolutionary significance for passerine diversification and suggests that programmed DNA elimination may be an overlooked yet phylogenetically widespread mechanism in many understudied animal lineages.

## Main

Multicellular organisms generally maintain the same genome in all of their cells. However, there are notable exceptions where programmed DNA elimination results in genomic differences between somatic and germline cells. This phenomenon has been observed across a broad range of eukaryotes, including lineages within various animal groups like nematodes, dipteran insects, cyclostome fishes, and songbirds (Wang and Davis 2014; Smith et al. 2021). One form of programmed DNA elimination is the germline-restricted chromosomes (GRCs), which are eliminated from the somatic cells early in embryogenesis and are only present in the germline cells. The first GRCs in birds were discovered in the zebra finch (Pigozzi and Solari 1998) and the Bengalese finch (Del Priore and Pigozzi 2014), and their presence has been confirmed in multiple songbird species since then (Torgasheva et al. 2019; Borodin et al. 2022). A striking finding was the substantial variation in GRC size, ranging from macro-GRCs exceeding 150 Mb to micro-GRCs as small as 5 Mb, with few intermediate forms. Interestingly, this variation does not align with phylogenetic relationships, as even closely related species can show dramatically different GRC sizes (Torgasheva et al. 2019; Borodin et al. 2022; Sotelo-Muñoz et al. 2022).

Passerines constitute the most diverse avian order, comprising around 6,700 out of 11,000 bird species, or roughly two thirds of all bird diversity (Gill et al. 2025). They emerged approximately 47 million years ago (Oliveros et al. 2019) and are divided into three major lineages: New Zealand Wrens, suboscines, and oscines (Infraorders Acanthisitti, Tyranni, Passeri, respectively, *sensu* Cracraft 2014). New Zealand Wrens are only composed of two extant species, while suboscines have ∼1,400 species and oscines, or songbirds, ∼5,300 species (Gill et al. 2025). Although much is known about passerine ecology, behaviour, physiology, and genetics, the presence of GRCs remained largely overlooked until recently. Still, the evolutionary history of the GRC—especially its occurrence beyond oscines and its potential role in germline development—remains poorly understood.

Analyses of a zebra finch macro-GRC draft assembly revealed a high enrichment of paralogous gene copies (Kinsella et al. 2019; Asalone et al. 2021), based on coverage ratio comparisons between liver and testis tissues, as well as single-nucleotide variants (SNVs). Comparable patterns in micro-GRC assemblies of blue tit (Mueller et al. 2023) and nightingales (Schlebusch et al. 2023), both of which belong to Passerides together with zebra finch (Oliveros et al. 2019) suggest that gene duplication onto the GRC is a recurring evolutionary phenomenon across this subgroup of oscines. These findings indicate that numerous genes have been incorporated into the GRC at different evolutionary timepoints, with a notable concentration of recent duplication events.

Here, we uncover the long-term evolutionary trajectories of the GRC by generating the first germline reference genomes for the zebra finch (*Taeniopygia guttata castanotis*) among oscines and the rusty-margined flycatcher (*Myiozetetes cayanensis*) among suboscines, using two complementary long-read sequencing technologies. Moreover, we analyze germline draft genomes from 23 oscines, two suboscines, and one parrot as an outgroup, together spanning all major lineages of passerine birds except Acanthisittidae. We uncover the GRC’s structural and functional evolution, gene retention patterns, and contribution to genomic innovation, underscoring the significant role of programmed DNA elimination in avian genome evolution.

### Rapid evolution of gene content and repeat content

We first aimed for an in-depth characterization of the composition of GRCs for comparison across species. To achieve this goal, we utilized assemblies based on long-read sequencing technologies. We selected the zebra finch, which is a model for GRC behaviour during meiosis and development (Itoh et al. 2009; Pei et al. 2022; Dedukh et al. 2025), and for the first time, a suboscine species—the rusty-margined flycatcher. We generated the first avian germline reference genomes using both accurate long-read PacBio HiFi sequencing and ultra-long Oxford Nanopore sequencing from testis samples (Supplementary Material). In both species, some assembly graphs are linear, including several that span from telomere to telomere. To identify GRC contigs, we detected testis-specific k-mers by comparing testis and soma sequencing libraries from the same individual, and selected contigs enriched in these k-mers. These contigs corresponded to the most complex non-linear graphs in the assembly (Supplementary Material). Additionally, we incorporated the recently assembled GRC of the blue tit *Cyanistes caeruleus* (Mueller et al. 2023), generated using PacBio HiFi long-read sequencing. This species represents another oscine clade but with a micro-GRC. Comparative analyses of the three species showed that, while the standard A chromosomes (autosomes and sex chromosomes) are highly collinear and syntenic across species (Figure 1A), the GRC shows a distinct evolutionary history (Figure 1B). The lack of synteny between species indicates rapid sequence turnover and lineage-specific sequence gains on GRC. However, the long-term retention of certain genes (Supplementary Material) points to selective pressures preserving functional elements.

**Figure 1.**
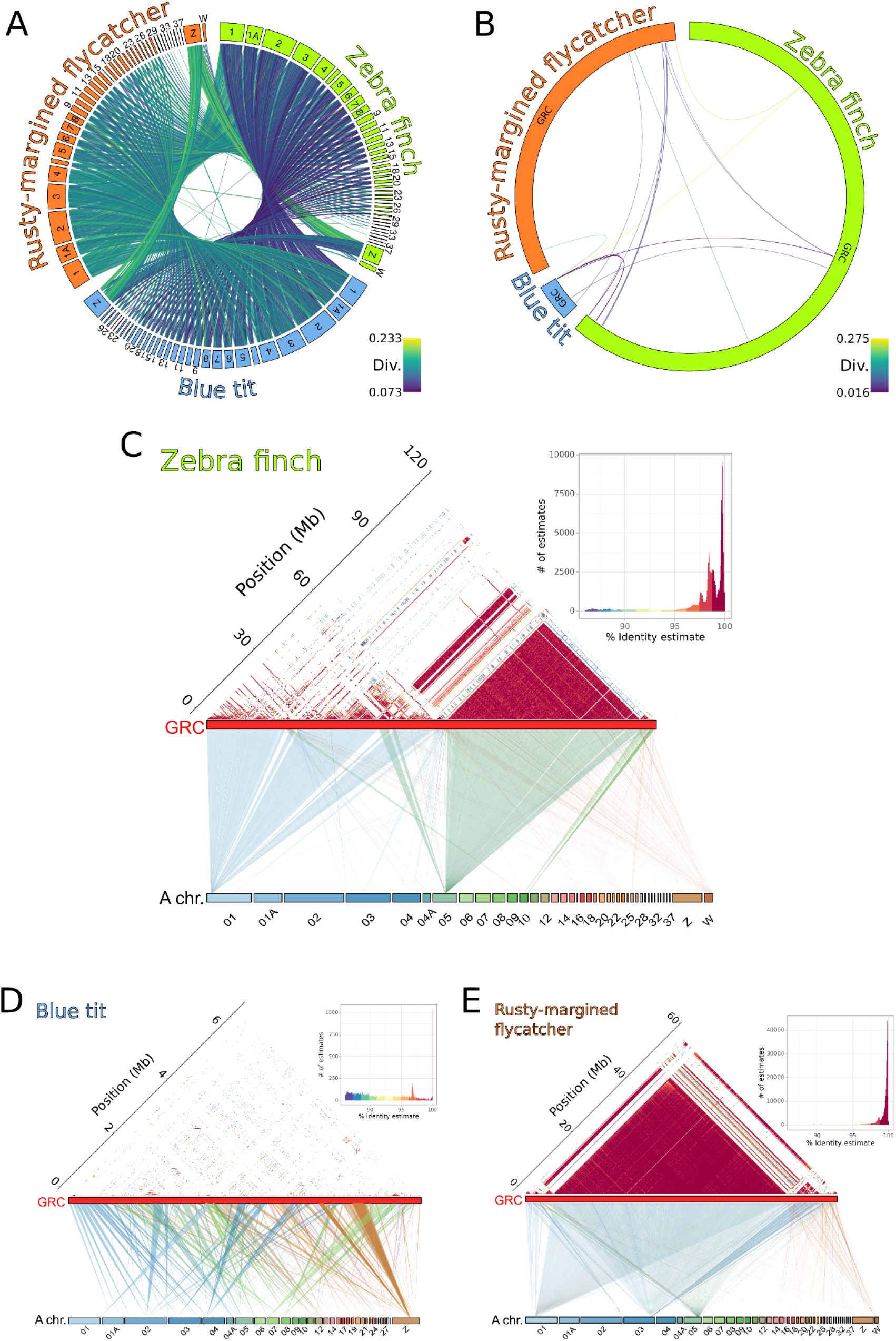
Extremely low synteny and complex organization of passerine GRCs. A) Circos plot of A chromosomes of the zebra finch (green), the blue tit (blue) and the rusty-margined flycatcher (orange) species shows high synteny and collinearity, reflecting conserved genome structure across passerines. B) Circos plots of their GRCs show minimal nucleotide sequence similarity, with few detectable connections between them, highlighting rapid divergence among GRCs. C-E) Dotplot heatmaps show pairwise nucleotide sequence similarity between concatenated GRC contigs of zebra finch (C), blue tit (D), and rusty-margined flycatcher (E). Red color indicates high identity, yellow intermediate identity, and blue low identity. Nucleotide alignments between the GRC and the A chromosomes from the same species are shown at the bottom of panels C-E, with one color for each A chromosome.

We aligned the GRC to the A chromosomes from the same germline genome at the nucleotide level. The zebra finch macro-GRC (∼160 Mb) is highly enriched in repetitive sequences, particularly with two major repeats derived from one single-copy region of chromosomes 1 and 5, respectively (Figure 1C). In contrast, the blue tit micro-GRC (∼7 Mb) predominantly consists of regions paralogous to most A chromosomes, but these are present in only a few copies on the GRC (Figure 1D). Meanwhile, the rusty-margined flycatcher meso-GRC (∼59 Mb) shows a similar pattern to the zebra finch with its main repeat derived from single-copy regions paralogous to chromosomes 1, 3 and 5 (Figure 1E).

### Functional Constraints and Retention of Key Genes

We identified 112, 145 and 125 protein-coding gene symbols in the GRC-linked regions of the rusty-margined flycatcher, blue tit and zebra finch, respectively, nearly all having paralogs on the A chromosomes and many having multiple copies on the GRC (Supplementary Material). To further investigate the gene content and evolutionary dynamics of GRCs across the Passeriformes phylogeny, we analyzed 10x Genomics Chromium linked-read libraries from somatic and germline tissues (testis) of 25 species representing key families of oscines: Corvides: Corvidae, Pachycephalidae, and Paradisaeidae; Passerides: Petroicidae, Hirundinidae, Paridae, Muscicapidae, Fringillidae, and Estrildidae; deep-branching oscines: Menuridae, Maluridae, and Meliphagidae); suboscines: Pittidae and Tyrannidae; and a parrot as an outgroup (Supplementary Material). No genes were universally present in all of the assemblies or read mappings (Figure 2) but, instead, hundreds of protein-coding gene symbols present only on the GRCs of one or two sampled families (Supplementary Material).

**Figure 2.**
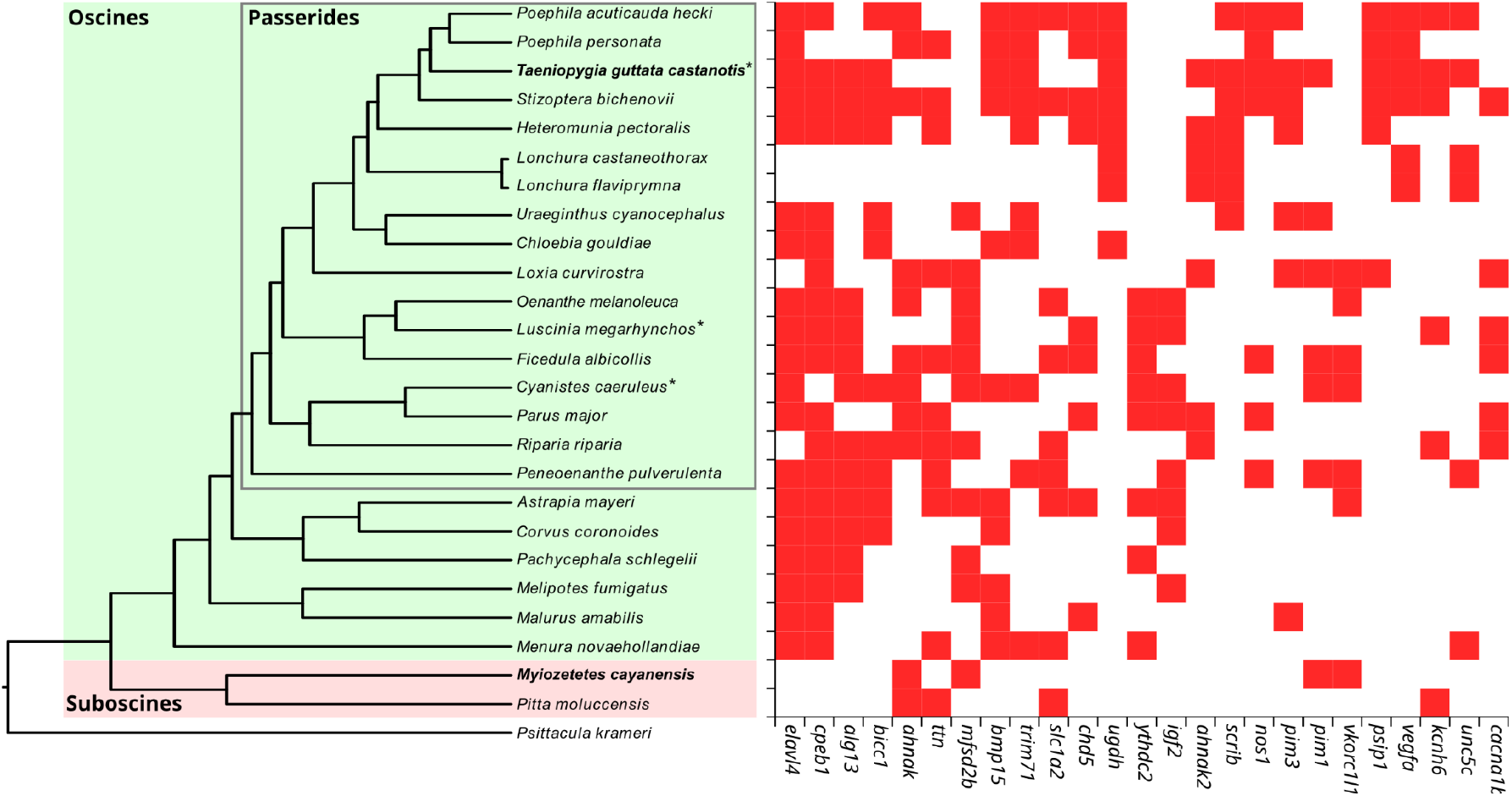
Dynamic evolution of GRC gene content across major passerine lineages. Left, maximum likelihood phylogeny of the 25 sampled passerines and one parrot outgroup, with testis and soma from the same individual sequenced with 10x Genomics linked reads (Supplementary Material). Bold species names indicate long-read germline assemblies and asterisks indicate previously generated linked-read datasets. Right, presence (red) of the 25 most widespread GRC-linked gene symbols (full table in Supplementary Material) based on the presence of testis-specific SNVs via mapping of 10x Genomics reads against the zebra finch transcriptome. Overrepresentation analysis of this subset of GRC-linked genes suggests an enrichment in biological processes such as intracellular neuronal signalling, mRNA stabilization, and activated cell proliferation (Supplementary Material).

Similarly, some GRCs were enriched in specific transposable elements, endogenous retroviruses, or satellite repeats, while others were less repetitive than the A-chromosomal background (Supplementary Material), suggesting rapid GRC sequence turnover across deep and recent evolutionary timescales.

We reconstructed the evolutionary history of a subset of phylogenetically widespread GRC genes by extracting their sequences from the A chromosomes and GRCs from both the draft germline and soma assemblies as well as the annotated reference germline assemblies (Supplementary Material). Our phylogenetic analyses confirmed that *cpeb1* was copied onto the GRC in the common ancestor of suboscines and oscines (Schlebusch et al. 2023), but is missing from the two suboscine GRCs analyzed (Figure 3A). In contrast, we detected the *pim1* gene on the GRCs of both zebra finch and rusty-margined flycatcher (Supplementary Material), providing strong evidence for a shared evolutionary origin of the GRC in early passerine diversification. We further demonstrated that genes can be independently copied, retained, or lost over time. For instance, *mfsd2b* was copied onto the GRC early in oscines but was acquired more recently in the rusty-margined flycatcher lineage within suboscines (Figure 3B).

**Figure 3.**
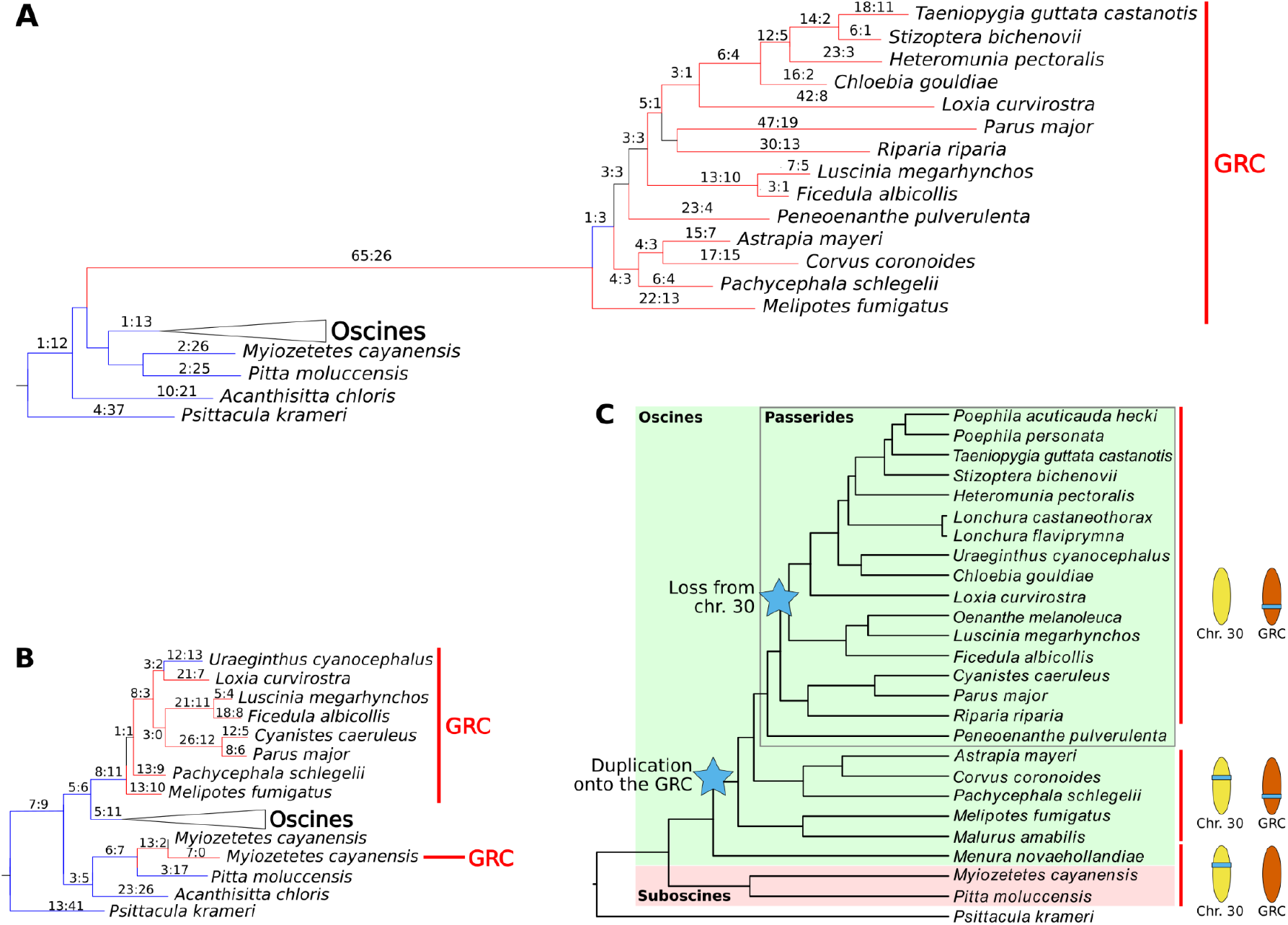
Complex evolutionary histories of GRC genes. A) *cpeb1* is an example of a GRC gene which arose from an ancient duplication onto the GRC in the common ancestor of oscines and suboscines. B) *mfsd2b* is an example of a GRC gene which underwent independent duplications onto the GRC with oscines and suboscines. C) *zglp1* is the only zebra finch GRC gene whose paralog was lost from the A chromosomes after duplication onto the GRC. The illustrated scenario is based on the *zglp1* gene tree and A-chromosomal synteny analyses of the germline reference genomes generated in this study (Supplementary Material). A-B) Maximum likelihood phylogenetic trees with 100,000 bootstrap replicates. Branch labels indicate the number of non-synonymous vs. synonymous substitutions, with red branches indicating positive selection and blue branches indicating purifying selection.

An even more extreme example is *bmp15*, which underwent a total of five independent duplication events across different lineages (Supplementary Material). Notably, the *zglp1* gene was duplicated onto the GRC in a common ancestor of most oscines, but the A-chromosomal copy is absent from the region between the *fdx2* and *icam5* genes on zebra finch chromosome 30, suggesting loss from the A chromosomes (Figure 3C, Supplementary Material). Importantly, several GRC genes show signatures of positive selection on the branches directly following their duplication, as indicated by an excess of non-synonymous substitutions (Figure 3A-B, Supplementary Material).

### Evolutionary and functional significance of GRCs

Our findings position GRCs as dynamic genomic innovators present in the majority of bird species, challenging the traditional classification as mere evolutionary anomalies. We show that GRCs are frequently reshaped over evolutionary timescales, rather than being genomic vestiges or repositories for surplus DNA (Johnson Pokorná and Reifová 2021). This stands in sharp contrast to the remarkable stability of avian A chromosomes in terms of synteny and gene content (Bravo et al 2021) and is even faster than the rapid evolution of avian W chromosomes (Peona et al. 2021; 2022). Thus, GRCs act as an evolutionary arena for gene innovation and conflict resolution (Vontzou et al. 2023), illuminating previously unrecognized aspects of the speed and mode of genome evolution in birds.

Our study provides the first direct evidence for the existence of GRCs in likely all ∼1,400 suboscine passerine species, broadening their known occurrence beyond all the ∼5,300 oscine passerines, i.e., songbirds. This discovery demonstrates their ubiquity within the Passeriformes and strongly supports a common ancestral origin predating the suboscine-oscine split, ∼44 million years ago (Oliveros et al. 2019). The sustained presence of GRCs since that split suggests they fulfill essential functional roles that have contributed to their stable evolutionary inheritance and developmental elimination despite extensive genomic reshaping and turnover across passerine lineages.

Our sampling of germline genomes across all major lineages of oscines and suboscines represents the first genomic study across different evolutionary timescales of any avian GRC system. The low degree of GRC synteny and extensive lineage-specific rearrangements across species reveal dynamic restructuring processes, sharply contrasting the highly conserved A chromosomes. Comparisons across recent evolutionary timescales show rapid evolutionary turnover (Kinsella et al. 2019; Sotelo-Muñoz et al. 2022; Schlebusch et al. 2023; Mueller et al. 2023), and our dense phylogenetic sampling highlights that this pattern can be generalized across passerine GRCs. We found that the GRC is under different dynamics of selective pressures or genetic drift compared to the A chromosomes, which we speculate to be driven by unique functional constraints or ongoing genomic conflict within the germline. These observations open intriguing questions regarding the specific mechanisms—such as cell cycle dynamics, recombination, meiotic drive, or selective gene retention—governing this remarkable rate of genomic evolution.

Despite extensive turnover, a subset of genes has been retained under purifying selection or positive selection (Figure 3, Supplementary Material; Biederman et al. 2018; Kinsella et al. 2019), suggesting functional relevance and non-random retention. The preservation of these genes strongly suggests their critical functional roles in germline development. A particularly notable discovery is the gene *pim1*, which is the only GRC gene retained by shared ancestry in both the zebra finch and the rusty-margined flycatcher germline reference genomes across 44 million years of divergence. This shared retention provides strong molecular evidence supporting a single origin of the GRC in the common ancestor of nearly all passerines, and highlights the deep evolutionary persistence of selected GRC genes. Even more compelling is the detection of positive selection in genes like *cpeb1* on the branch leading to the GRC ancestor, indicating adaptive evolution of the GRC paralog. Notably, *cpeb1* is absent from all sampled suboscines, including the germline reference genome of the rusty-margined flycatcher, suggesting that the gene was lost early in that lineage. Its absence from the PacBio long-read germline assembly of the blue tit, but presence in the great tit draft germline assembly exemplifies a relatively recent loss of *cpeb1*. These patterns provide the important insight that even genes retained for tens of millions of years can eventually be lost, reflecting a dynamic balance between functional importance and rapid evolutionary change.

A particularly striking example of the dynamic functional capacity of the GRC is demonstrated by the gene *zglp1*. Originally located on chromosome 30, *zglp1* was subsequently duplicated onto the GRC early during oscine diversification and then lost from the autosome in the common ancestor of most Passerides, a group comprising ∼4,000 oscine species (Gill et al. 2025), making *zglp1* the first known case of a truly germline-specific gene with no remaining copy on the A chromosomes and thus complete absence from somatic cells. Given the well-established role of *zglp1* in female germ cell development and fertility in mice (Nagaoka et al. 2020), its presence on the GRC may have permitted or buffered the loss of the A-chromosomal copy, underscoring the potential of the GRC to retain essential germline functions. The event suggests that the GRC can act as a genomic reservoir, capable of safeguarding essential gene functions within the germline. We hypothesize that in nearly all of the 4,000 Passerides species, the essentiality of *zglp1* has rendered the GRC indispensable, preventing its loss due to the absence of a compensatory A-chromosomal *zglp1* copy. Furthermore, this may reflect a case of minimizing antagonistic pleiotropy, whereby restricting gene presence to the germline avoids potential deleterious effects of misexpression in somatic tissues (reviewed in Smith et al. 2021 and Vontzou et al. 2023). The germline limitation of *zglp1* is thus a compelling illustration of adaptive gene content evolution, reflecting the dynamic interplay between gene loss from the A chromosomes and functional maintenance through the GRC. Further investigation into similar cases across passerines may illuminate broader mechanisms of gene duplication and adaptation associated with GRC evolution. The presence of such distinctive gene content likely contributes significantly to the evolutionary persistence of GRCs, preventing their selective loss or degradation. Investigating the specific biological functions of these genes within the germline is a promising avenue to further understand their selective advantages and evolutionary maintenance.

To conclude, our analyses reveal the GRC to be a key genomic innovator involved in passerine genome evolution, highlighting their capacity for functional innovation, adaptation, and intragenomic conflict resolution. These results also contribute to a broader understanding of programmed DNA elimination across distant taxa. In lampreys, GRCs contain hundreds of expressed genes involved in germline development and cancer-related pathways, many of which arose from repeated duplication events, both from A chromosomes and on the GRC (Smith et al. 2018; Timoshevskaya et al. 2023). Hagfishes show similar patterns of duplication and expansion on their GRCs (Marlétaz et al. 2024). By contrast, in fungus gnats, GRCs contain paralogs of thousands of different genes, with little evidence of gene degradation, suggesting a different evolutionary trajectory (Hodson et al. 2022). These cross-lineage comparisons reveal that GRCs, while independently evolved, may share a convergent role as platforms for germline-specific genomic innovation and specialization. Future work should clarify novel functions of GRC gene paralogs and the evolutionary forces shaping their retention and diversity.

## Methods

### Genome assembly and identification of GRC contigs

We generated testis assemblies from long-read libraries sequenced on the Pacific Biosciences HiFi and Oxford Nanopore Technologies platforms obtained from a single individual both the zebra finch and the rusty-margined flycatcher (Supplementary Table 1, Supplementary Material), using hifiasm v0.19.5 (Cheng et al. 2024) with the option “-l0” to disable the purge duplication step. We retained the primary assembly for downstream analyses. To identify GRC contigs in these two assemblies, we first counted k-mers (k=21) with Meryl v1.4.1 (Rhie et al. 2020) in the testis PacBio HiFi reads and the soma Illumina reads from the same individual. We then compared the k-mer sets to identify testis- and soma-specific k-mers with the “trio/hapmers.sh” script from the Merqury suite v.1.3 (Rhie et al. 2020). The soma-specific k-mers were used as a negative control. We quantified the occurrences of testis- and soma-specific k-mers in the assemblies with the meryl-lookup (Rhie et al. 2020) and normalized these counts by contig length. We defined as GRC-linked those contigs where at least 3% of the k-mers were testis-specific. Below this k-mer density, we observed a sharp signal drop-off (Supplementary Material). Additionally, we generated 10x Genomics Chromium linked-read libraries from testis and soma samples originating from one individual each for 25 bird species (Supplementary Table 1, Supplementary Material) with Supernova2 (Weisenfeld et al. 2017) using the pseudohap option.

### Genome annotation

We designed a custom pipeline to annotate genes present in the GRC contigs of our zebra finch and rusty-margined flycatcher assemblies (Supplementary Material), as well as for the blue tit (Mueller et al. 2023, NCBI Bioproject PRJNA925103). We performed homology-based annotation with GeMoMa v1.9 (Keilwagen et al. 2019) and *de-novo* annotation with the deep-learning annotation tool Helixer v0.3.4 (Holst et al. 2023) through its web server (https://www.plabipd.de/helixer_main.html) with the model for Vertebrates. Also, we identified pseudogenes using the PseudogenePipeline.py v2.0.0 script (https://github.com/ShiuLab/PseudogenePipeline). We converted all the resulting GFF or GTF files into standardized GFF format using AGAT (Dainat et al. 2019). We then merged the three annotation sources hierarchically, prioritizing GeMoMa, followed by Helixer, and then the Pseudogene pipeline annotations in case of overlap.

Afterwards, we extracted the protein sequences with AGAT and used DIAMOND v2.1.9 (Buchfink et al. 2021) for local alignment against the NCBI non-redundant protein database. We retained the top match with a gene symbol as the gene name.

### Genome alignments

The GRC reference assemblies of zebra finch and rusty-margined flycatcher consisted of a few hundred contigs, while the blue tit GRC had a few dozen. For simplified visualization, we aligned the GRC contigs of the zebra finch, blue tit, and rusty-margined flycatcher against the A chromosome from the same species with D-GENIES v1.5.0 (Cabanettes and Klopp 2018) through its web server (https://dgenies.toulouse.inra.fr/run). We then ordered the GRC contigs based on their similarity and concatenated them. We aligned the A chromosomes and GRCs of the three species to each other, respectively, using minimap2 v2.25 (Li 2018) with default options. We used the R package circlize v0.4.14 (Gu et al. 2014) to visualize these alignments as Circos plots. We also used minimap2 to align the GRC to the A chromosomes from the same species. In this case, we used SyntenyPlotteR v1.0.0 (Quigley et al. 2023) for visualization. We also generated heatmap dotplots for the concatenated GRC contigs with ModDotPlot v0.9.4 (Sweeten et al. 2024).

### SNV calling for testis

We used the 10x Genomics linked-read libraries from testis and soma of each of the 25 species to map against the zebra finch taeGut2 transcriptome, after removing redundant sequences, mainly from different isoforms, with CD-HIT-EST (Fu et al. 2012) using the options “-M 0 -aS 0.8 -c 0.8 -G 0 -g 1”. These settings clustered sequences with 80% similarity across 80% of their transcript length, using local alignment and a greedy algorithm, as in Kinsella et al. (2019). We next removed linked-read barcodes with Longranger basic (10x Genomics) and performed adapter and quality trimming with Trimmomatic v0.36 (Bolger et al. 2014) using the options “PE -phred33 ILLUMINACLIP: 2:1:10 SLIDINGWINDOW:4:20 MINLEN:20”. We used the resulting trimmed reads to perform SNV calling using the whatGene pipeline (https://github.com/fjruizruano/whatGene) as described in Ruiz-Ruano et al. (2019) and Kinsella et al. (2019). Thus, we generated a list of candidate genes for the GRC of each species and summarized these results in a presence/absence matrix (Supplementary Material). We visualized this matrix together with a maximum likelihood tree built with RAxML v8.2.12 (Stamatakis 2014), using mitogenomes assembled with MITObim v1.9.1 (Hahn et al. 2013) from all the 10x Genomics linked-read libraries. We observed congruence of this tree with Oliveros et al. (2019) and Pei et al. (2022).

### GRC gene phylogenies

We extracted gene sequences from the Supernova2 assemblies for a subset of genes present among most of the 25 sampled species (Supplementary Material). For these genes, we retrieved the coding DNA sequences and amino acid sequences from GenBank for the relevant species (Supplementary Material). We used these sequences as references in the CAPTUS software (Ortiz et al. 2023). Next, we aligned the sequences with MAFFT v7.453 (Katoh and Standley 2013) using the “linsi” options and manually inspected the alignments with Geneious v4.8.5 (Drummond et al. 2009) to remove uninformative sequences. We then realigned the filtered sequences using GUIDANCE2 v2.0.2 (Sela et al. 2015) with PRANK v.121211 (Löytynoja 2014), applying 100 bootstrap replicates to discard unreliable regions with scores below 0.93. After this, we refined the alignments by removing codons represented in fewer than 70% of sequences, and sequences with information in fewer than 50% of aligned sites. We used the resulting alignments to construct maximum likelihood phylogenetic trees with IQ-TREE v.2.2.2.6 (Minh et al. 2020), using 10,000 ultrafast bootstrap replicates. We defined as GRC lineages those composed of sequences exclusively present in testis libraries (Supplementary Material). When multiple near-identical GRC-linked or A-chromosomal sequences were detected for the same species, we kept a single sequence to simplify tree visualization and reduce tree reconstruction bias. We used the free-ratio model from codeml, part of the PAML v4.10 suite (Yang 2007), to estimate the number of nonsynonymous and synonymous substitutions for each branch. We minimized the potential overestimation of this model by excluding sites with ambiguous bases or alignment gaps. Finally, we visualized and edited the trees with FigTree v.1.4.5 (Rambaut 2018).

## Data availability

Prior to journal submission of this manuscript, we will submit all PacBio HiFi, Oxford Nanopore, and 10x Genomics Chromium and Illumina libraries as well as all assemblies to the relevant databases (Bioproject PRJNAXXXX). Custom code will be made available on GitHub (https://github.com/fjruizruano).

## Acknowledgements

We thank Juan Pedro M. Camacho for inspiring some of the ideas explored in this work. We are grateful to the Suh Lab, the GRC Brainstorming group, and Erich Jarvis for helpful discussions throughout this six-year project. We thank Maria Luisa da Silva (Laboratory of Ornithology and Bioacoustics, Federal University of Para, Belem, Brazil) for her contribution and assistance in collecting rusty-margined flycatcher samples, Katrin Martin and Moritz Hertel (Department of Ornithology, Max Planck Institute for Biological Intelligence, Seewiesen, Germany) for their assistance in collecting zebra finch samples, and Jan Engler, Antje Bakker, Viki Vandomme, and Diederik F. Strubbe (Terrestrial Ecology Unit, Department of Biology, Ghent University, Ghent, Belgium) for providing ring-necked parakeet samples. This work was funded by a Consolidator Grant of the European Research Council (101002158 GermlineChrom) to AS, a Future Research Leaders Grant of the Swedish Research Council Formas (2017-01597) to AS, a Project Grant of the Swedish Research Council Vetenskapsrådet (2020-04436) to AS and FJRR, a Nilsson Ehle Endowment of the Royal Physiographic Society of Lund to AS, a Marie Skłodowska-Curie Individual Fellowship (875732 birdGRC) to FJRR, a Bioinformatics Long-term Support project of the National Bioinformatics Infrastructure Sweden to AS and FJRR, a Postdoctoral fellowship from Sven och Lilly Lawskis fond (Sweden) to FJRR, a Starting Grant of the Swedish Research Council Vetenskapsrådet (2020-03866) to OMPG, funds from the Max Planck Society to BK, a Project Grant of the Czech Science Foundation (23-07287S to RR and TA, 25-17195S to RR), a Project Grant of the National Institutes of Health (K99-HG014014-01) to MTB, and a Project Grant of the Ministry of Science and Higher Education of the Russian Federation (#FSUS-2024-0018) to LM. The computations were enabled by resources provided by the National Academic Infrastructure for Supercomputing in Sweden (NAISS) at UPPMAX, funded by the Swedish Research Council through grant agreement no. 2022-06725.

## Notes

### Competing Interest Statement

The authors have declared no competing interest.

